# Berberrubine Inhibits *Helicobacter pylori* by Inducing Oxidative Stress

**DOI:** 10.1101/2025.03.06.641886

**Authors:** Min-Zhi Jiang, Chang-Yu Wang, Kai Wang, Yu Jiang, Xue Wang, Xin-Chi Feng, Shi-Jie Cao, Li-Qin Ding, Shuang-Yu Bi, Feng Qiu, Shuang-Jiang Liu, Chang Liu

**Author notes:** **Corresponding author** Correspondence to Feng Qiu, Chang Liu and Shuang-Jiang Liu.

## Abstract

*Helicobacter pylori* is a prevalent pathogen that colonizes the pylorus and contributes to the pathogenesis of gastric cancer. The increasing antibiotic resistance in *H. pylori* presents a substantial challenge for effective eradication, making the development of more effective therapeutic strategies imperative. In this study, we screened 37 pure compounds (purity >95%), 3 ethanol extracts, and 6 aqueous extracts from 6 different medicinal herbs for their potential inhibitory effects against *H. pylori in vitro*. Among the tested compounds, 15 exhibited inhibitory activity, with berberrubine demonstrating the strongest effect, characterized by a minimum inhibitory concentration (MIC) of 0.011 mg/ml. The comparative transcriptome analysis revealed a significant upregulation of the oxidative stress response related genes, such as *fumC* and *sodB*, in *H. pylori treated with berberrubine*. Further *in vitro* assays confirmed that berberrubine exerted antibacterial activity by stimulating oxidative stress, as evidenced by increased reactive oxygen species (ROS) production and impaired cell membrane integrity. Additionally, similar inhibitory tests were applied to *Escherichia coli*, a common gut opportunistic pathogen. The results showed that berberrubine inhibited *E. coli* through the same mechanism observed for *H. pylori*. In conclusion, berberrubine is a potent antibacterial agent against gastrointestinal pathogens, including *H. pylori* and *E. coli*, making it a promising candidate for further development as an antimicrobial therapeutic

**Importance:** This study identified berberrubine as a potent antibacterial agent against gastrointestinal pathogens, including *Helicobacter pylori and Escherichia coli* by inducing cellular oxidative stress. Through increased reactive oxygen species (ROS) production and disruption of bacterial cell membrane integrity, berberrubine demonstrated strong efficacy in inhibiting drug-resistant pathogens. These findings highlight its potential as a promising therapeutic candidate for eradicating drug-resistant gastrointestinal pathogens.

## Introduction

Increasing evidence has shown that gastrointestinal microbes contributed to the development of various cancers (1-5). *Helicobacter pylori* is a common pathogenic bacterium colonizing the human gastric mucosa, particularly the pyloric region, and is directly associated with the occurrence and progression of gastric cancer (4, 6). In the absence of effective drug intervention, *H. pylori* can persist in the human body for decades, contributing to chronic inflammation and cellular damage. Alarmingly, approximately 600,000 new cases of gastric cancer reported annually are related to *H. pylori* infection, prompting the World Health Organization (WHO) to classify it as a Class I carcinogen (7). The persistence of *H. pylori* is largely attributed to its remarkable resistance to various antibiotics, posing a significant challenge to its effective eradication. The most commonly-used treatments for *H. pylori* infections in clinicals are quadruple therapies. These regimens are either bismuth-based, involving a combination of bismuth (a topical agent), tetracycline, metronidazole, and a proton pump inhibitor (PPI), or non-bismuth quadruple therapy, also known as concomitant therapy (8, 9). The latter combines a PPI with amoxicillin, clarithromycin, and metronidazole, administered simultaneously rather than sequentially (10). Although these strategies largely improve treatment efficacy and reduce the risk of antibiotic resistance, challenges persist due to the growing prevalence of drug resistance and frequent treatment failures. Therefore, it is imperative to explore more effective bactericidal strategies against *H. pylori* as alternatives or complements to existing antibiotic therapies.

Increasing number of research has demonstrated that plant-derived natural compounds exhibit inhibitory effects against pathogens including *H. pylori* (11-14). Traditional plant extraction techniques, including alcohol and aqueous extraction, yield crude extracts with broad-spectrum components, which exhibit inhibitory activity against *H. pylori*. For example, methanol extracts from *Carum carvi* (seed), *Elettaria cardamomum* (seed), *Gentiana lutea* (roots), *Juniper communis* (berry), *Lavandula angustifolia* (flowers), *Melissa officinalis* (leaves), *Mentha piperita* (leaves), *Pimpinella anisum* (seed), *Myroxylon peruiferum* and *Terminalia macroptera* (root) have been reported to exert inhibitory activity against *H. pylori* (15, 16, 17). Although these crude extracts exhibit antimicrobial properties, their complex nature limits their use by complicating toxicological evaluations and formulation development, particularly due to uncertainties in drug metabolism and pharmacokinetics (17-24). Therefore, in recent years, research paradigms have gradually shifted towards the isolation, purification, and structural optimization of single active compounds, such as Cinnamic acid (MIC= 80-200 μg/mL), Paeonol, benzoic acid, methyl gallate (MIC= 80-320 μg/mL), 1,2,3,4,6-Penta-O-galloyl-β-D-glucopyranose (MIC= 320-1280 μg/mL), and Chlorogenic acid (MIC= 312.5-1250 μg/mL) (13, 25-27). Among the many plant-derived antimicrobial compounds, the research on the isoquinoline alkaloid berberine has been the most representative (28, 29). This compound is primarily isolated from medicinal plants such as *Coptis chinensis* and *Phellodendron amurense*, and its unique tetracyclic structure gives it broad-spectrum antibacterial properties (29). Existing evidence confirms that berberine is one of the most effective plant-derived anti-*H. pylori* agent, with a MIC as low as 50 μg/mL, making it a strong alternative to antibiotics. Mechanistically, it exerts its antibacterial effect by increasing oxidative stress (30). Notably, clinical trials have demonstrated that berberine containing quadruple therapy is was not inferior to standard bismuth quadruple therapy in *H. pylori* eradication (31). These promising results suggest that plant-derived natural products could be a treasure trove of resources for discovering antibiotic alternatives, warranting further exploration.

In this study, by screening of 37 chemicals, 3 ethanol extracts, and 6 aqueous extracts isolated from 6 herbs on their inhibitory effects against *H. pylori*, we identified 12 antibacterial compounds. Berberrubine exhibited the strongest antibacterial activity (MIC= 11 μg/mL) through inducing oxidative stress and membrane damage in treated *H. pylori* cells. Validation with *E. coli* confirmed similar antimicrobial effects, suggesting its broader applicability as an antimicrobial agent. These findings provide a basis for further development of natural therapeutic for combating antibiotic-resistant *H. pylori* and other gastrointestinal pathogens.

## Results

### *In vitro* activity screening of herb extracts revealed that berberrubine exerted strong antimicrobial effects against *Helicobacter pylori*

To identify natural products with inhibitory effects on *H. pylori*, a total of 37 pure compounds (purity >95%), 3 ethanol extracts, and 6 aqueous extracts from herbs were tested *in vitro* for their MICs against *H. pylori*. As shown in Table1, 15 compounds from 4 herbs exhibited antibacterial activity, while the other 31 showed no activity. Notably, all 6 purified compounds and aqueous extracts derived from *Coptidis rhizoma* demonstrated inhibitory effects against *H. pylori*, with berberrubine showing the strongest antibacterial activity, achieving a minimum inhibitory concentration (MIC) of just 0.011 mg/ml. Additionally, two other compounds derived from *Coptidis rhizoma*, epiberberine and berberine, exhibited the second and third strongest antibacterial effects, with MIC values of 0.013 mg/ml and 0.025 mg/ml, respectively.

**Table 1.**
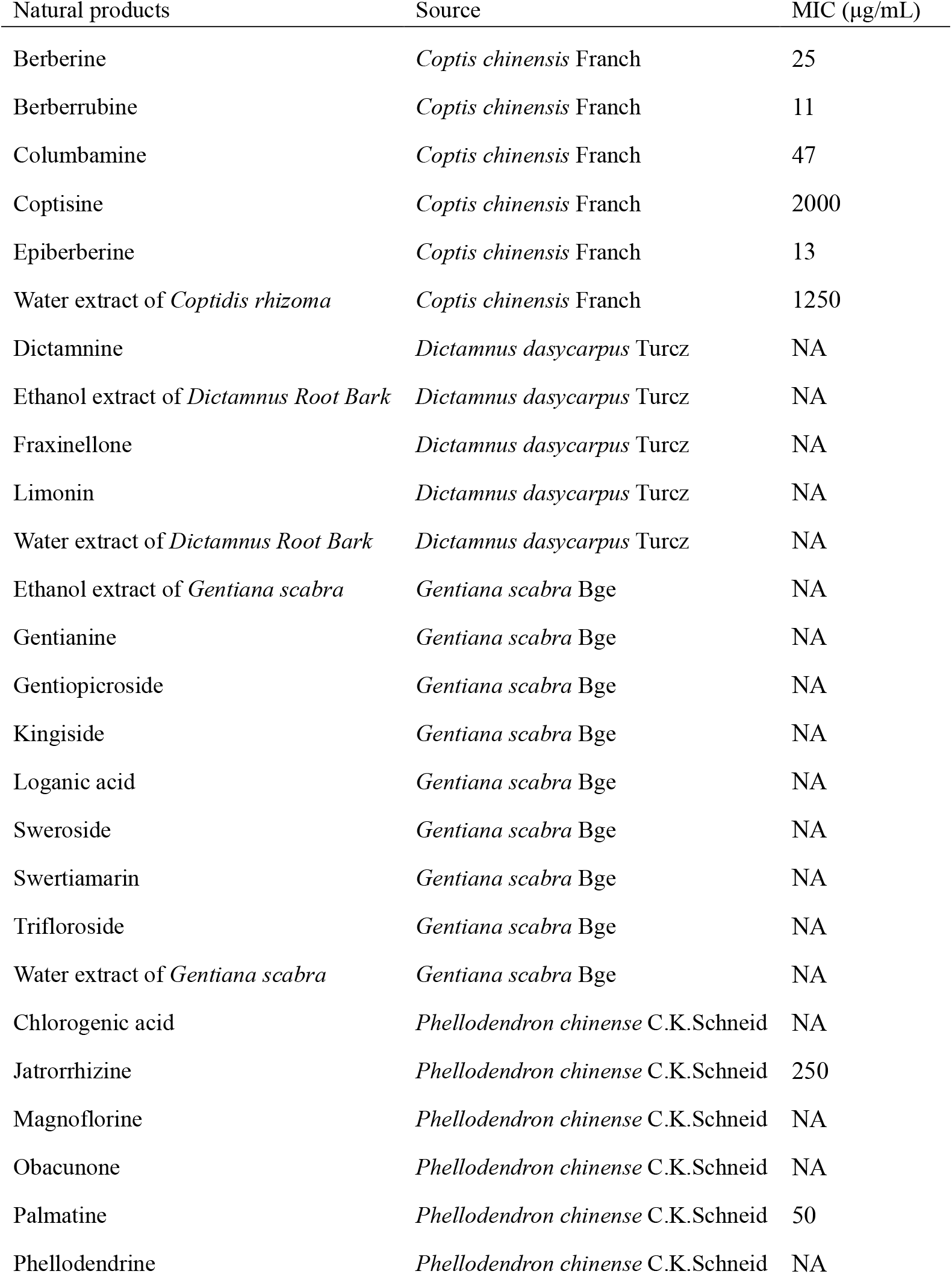
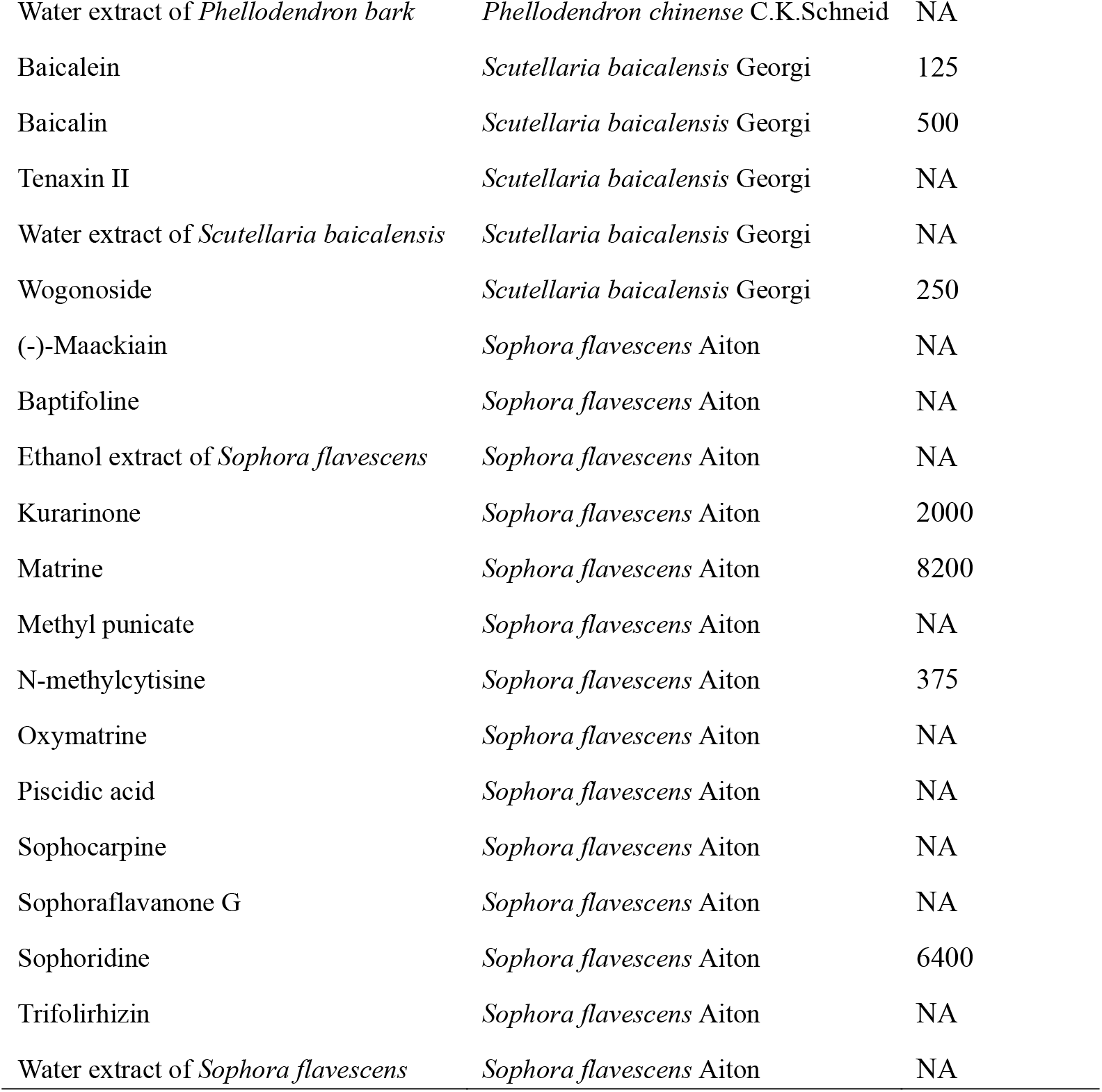
The minimum inhibitory concentration (MIC) of 46 natural extracts against *Helicobacter pylori*.

| Natural products | Source | MIC (µg/mL) |
| --- | --- | --- |
| Berberine | *Coptis chinensis* Franch | 25 |
| Berberrubine | *Coptis chinensis* Franch | 11 |
| Columbamine | *Coptis chinensis* Franch | 47 |
| Coptisine | *Coptis chinensis* Franch | 2000 |
| Epiberberine | *Coptis chinensis* Franch | 13 |
| Water extract of *Coptidis rhizoma* | *Coptis chinensis* Franch | 1250 |
| Dictamnine | *Dictamnus dasycarpus* Turcz | NA |
| Ethanol extract of *Dictamnus Root Bark* | *Dictamnus dasycarpus* Turcz | NA |
| Fraxinellone | *Dictamnus dasycarpus* Turcz | NA |
| Limonin | *Dictamnus dasycarpus* Turcz | NA |
| Water extract of *Dictamnus Root Bark* | *Dictamnus dasycarpus* Turcz | NA |
| Ethanol extract of *Gentiana scabra* | *Gentiana scabra* Bge | NA |
| Gentianine | *Gentiana scabra* Bge | NA |
| Gentiopicroside | *Gentiana scabra* Bge | NA |
| Kingiside | *Gentiana scabra* Bge | NA |
| Loganic acid | *Gentiana scabra* Bge | NA |
| Sweroside | *Gentiana scabra* Bge | NA |
| Swertiamarin | *Gentiana scabra* Bge | NA |
| Trifloroside | *Gentiana scabra* Bge | NA |
| Water extract of *Gentiana scabra* | *Gentiana scabra* Bge | NA |
| Chlorogenic acid | *Phellodendron chinense* C.K.Schneid | NA |
| Jatrorrhizine | *Phellodendron chinense* C.K.Schneid | 250 |
| Magnoflorine | *Phellodendron chinense* C.K.Schneid | NA |
| Obacunone | *Phellodendron chinense* C.K.Schneid | NA |
| Palmatine | *Phellodendron chinense* C.K.Schneid | 50 |
| Phellodendrine | *Phellodendron chinense* C.K.Schneid | NA |
| Water extract of *Phellodendron bark* | *Phellodendron chinense* C.K.Schneid | NA |
| Baicalein | *Scutellaria baicalensis* Georgi | 125 |
| Baicalin | *Scutellaria baicalensis* Georgi | 500 |
| Tenaxin II | *Scutellaria baicalensis* Georgi | NA |
| Water extract of *Scutellaria baicalensis* | *Scutellaria baicalensis* Georgi | NA |
| Wogonoside | *Scutellaria baicalensis* Georgi | 250 |
| (-)-Maackiain | *Sophora flavescens* Aiton | NA |
| Baptifoline | *Sophora flavescens* Aiton | NA |
| Ethanol extract of *Sophora flavescens* | *Sophora flavescens* Aiton | NA |
| Kurarinone | *Sophora flavescens* Aiton | 2000 |
| Matrine | *Sophora flavescens* Aiton | 8200 |
| Methyl punicate | *Sophora flavescens* Aiton | NA |
| N-methylcytisine | *Sophora flavescens* Aiton | 375 |
| Oxymatrine | *Sophora flavescens* Aiton | NA |
| Piscidic acid | *Sophora flavescens* Aiton | NA |
| Sophocarpine | *Sophora flavescens* Aiton | NA |
| Sophoraflavanone G | *Sophora flavescens* Aiton | NA |
| Sophoridine | *Sophora flavescens* Aiton | 6400 |
| Trifolirhizin | *Sophora flavescens* Aiton | NA |
| Water extract of *Sophora flavescens* | *Sophora flavescens* Aiton | NA |

### Berberrubine treatment upregulated genes related to oxidative stress in *Helicobacter pylori*

To elucidate the antibacterial mechanism underpinned the observed inhibitory activity, *H. pylori* cells were treated with berberrubine (TREAT in Fig. 1) and without (CK Fig. 1) for 2 hours, respectively. RNA-seq based transcriptomic analysis was conducted on bacterial cells at the initial time point (CK0 and TREAT0 in Fig. 1) and after 2-hour treatment (CK1 and TREAT1 in Fig. 1) to compare differential gene expressions. The Kendall correlation coefficients were employed to assess the similarity in transcriptional profiles between any two datasets, which indicated that in addition to drug treatment, cultivation time also affected gene expressions (Fig. 1A). The Principal Component Analysis (PCA) further revealed significant changes in the transcriptional levels between the CK0 to CK1 and TREAT0 to TREAT1 samples (Fig. 1B), which corroborated the influence of cultivation time on gene expressions. These results indicated that when analyzing differentially expressed genes (DEGs) caused by drug treatment, the transcriptional changes with time should be excluded. Subsequently, DEG analysis revealed that, compared to CK0, 9 genes were significantly upregulated and 29 genes were significantly downregulated in CK1. Similarly, in TREAT1 compared to TREAT0, 18 genes were significantly upregulated and 9 genes were significantly downregulated. To depict the overall response of *H. pylori* to berberrubine treatment and cultivation time, we performed functional annotation of all DEGs based on Gene Ontology (GO). Compared to CK0, CK1 mainly showed differences in molecular functions related to nucleoside and ribonucleoside binding and cell proliferation, indicating that cultivation time primarily affected cell growth and normal physiological activities. However, after berberrubine treatment, the most significantly enriched processes were related to oxidation reduction, energy metabolism, cellular respiration and drug metabolism, indicating that berberrubine enhances oxidative stress responses and cellular repair in *H. pylori* (Fig. 1E, 1F). These findings suggested that berberrubine stimulated oxidative stress responses and cellular metabolism in *H. pylori*, potentially helping the bacteria withstand cellular damage.

**Fig. 1:**
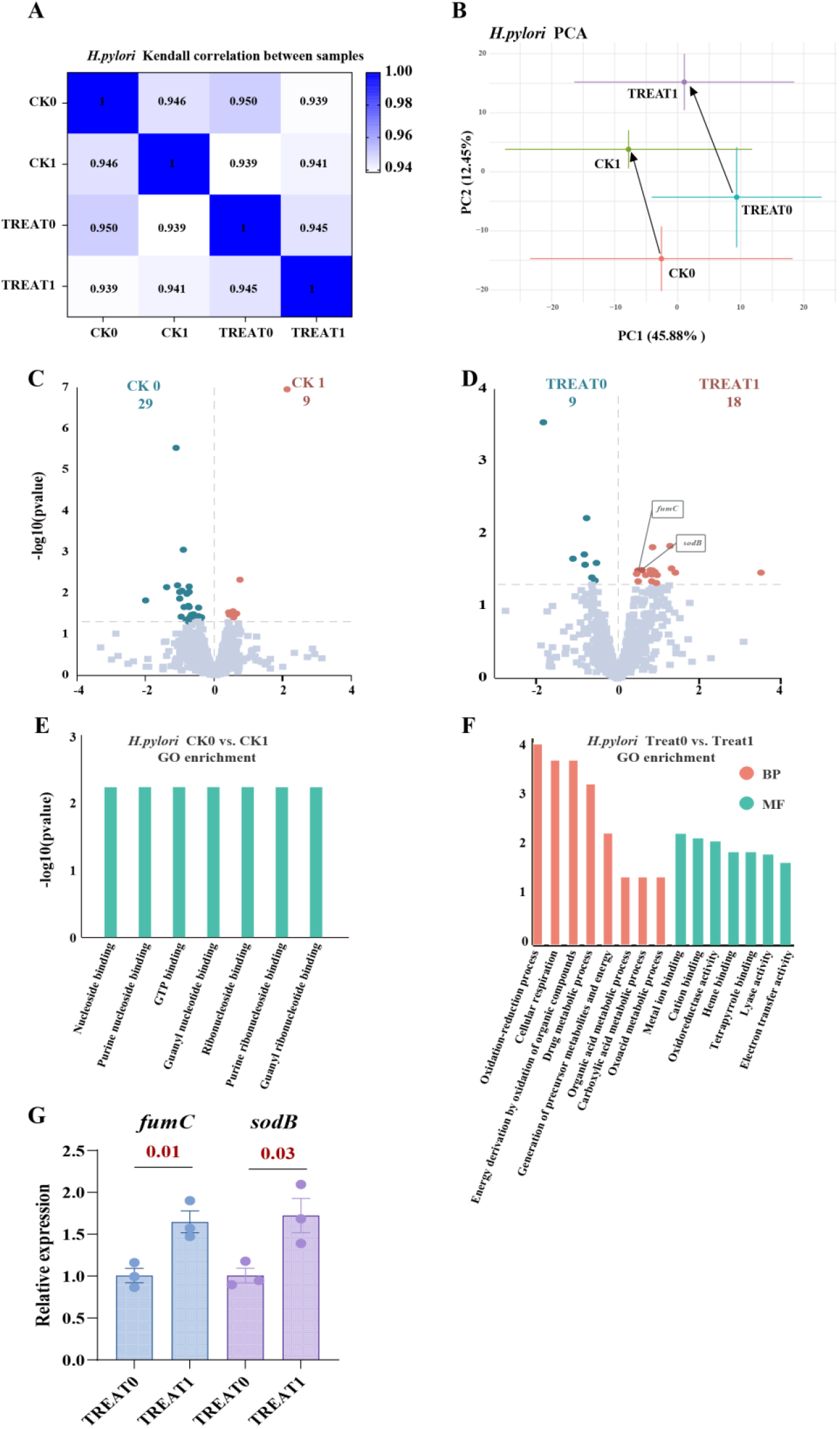
Transcriptomic analysis of *H. pylori* transcriptional changes after 2 h of berberrubine treatment. (A) Kendall correlation coefficient analysis comparing the gene expression levels between the CK and TREAT groups. (B) PCA analysis showing transcriptional changes between the CK0 to CK1 and TREAT0 to TREAT1 samples. (C) Volcano plot of differential genes for CK0 vs. CK1, with |log2FC| > 0 and p-value < 0.05. (D) Volcano plot of differential genes for TREAT0 vs. TREAT1, with |log2FC| > 0 and p-value < 0.05. (E) GO enrichment analysis results for the control group CK0 vs. CK1. (F) GO enrichment analysis results for the treatment group TREAT0 vs. TREAT1. (G) qPCR relative quantification of the *fumC* and *sodB* gene transcription level in treated group of *H. pylori*. Data were shown as the mean±S.E.M. The p value was calculated by Student’s t-test.

Among 18 genes upregulated in TREAT1 compared to TREAT0, three were also upregulated in CK1 relative to CK0, suggesting a potential influence from cultivation time (Table S1). Of the remaining 15 genes specifically upregulated by berberrubine treatment, six were functionally annotated as being associated with oxidative stress responses. These including *fumC*, encoding class II fumarate hydratase (OW489_RS06375/), *sodB*, encoding superoxide dismutase (OW489_RS07685), a pyruvate flavodoxin oxidoreductase (OW489_RS07460), a putative cytochrome-c peroxidase (OW489_RS00335), a putative cytochrome-b (Novel00152) and a putative iron manganese superoxide dismutase (Novel00227), all highlighted in red in Table S1. Notably, *fumC* and *sodB* have been previously reported to be responsive to oxidative stress. *fumC* catalyzed the conversion of fumarate to malate, thereby enhancing cellular energy metabolism and providing energy for cellular repair (32, 33), while *sodB* neutralized superoxide radicals, thereby protecting microbial cells from oxidative damage and maintaining cellular homeostasis (34). To verify these findings from RNA-seq, RT-qPCR was performed to assess the expression levels of *fumC* and *sodB*. Following berberrubine treatment of *H. pylori* cells under the same conditions as the RNA-seq experiment, the mRNA levels of both *fumC* and *sodB* were significantly increased (Fig. 1G), suggesting their involvement in a cellular defense mechanism against oxidative stress. Based on above evidence, we therefore inferred that berberrubine might exert its antibacterial effect by inducing oxidative stress and causing cellular damage.

### Berberrubine exerted antibacterial effects by increasing intracellular ROS levels and impairing cellular integrity

To experimentally verify the effect of berberrubine on oxidative stress in *H. pylori*, 2’,7’-dichlorofluorescein diacetate (DCFH-DA) was used to detect reactive oxygen species (ROS) in cells treated with 2× and 5× MIC concentrations of berberrubine. The results showed that ROS rapidly accumulated within 2 hours of treatment with both concentrations, and after 8 hours, ROS levels in the 5× MIC treatment group remained significantly elevated (Fig. 2A). These findings confirmed that berberrubine treatment led to a significant, dose-dependent increase in intracellular ROS levels. Furthermore, scanning electron microscopy (SEM) was used to observe potential changes in bacterial morphology caused by berberrubine treatment. In the control (CK) group, *H. pylori* cells appeared structurally intact, with clear, flat, smooth cell membrane boundaries, a full morphology, and homogeneous cytoplasm (Fig. 2B). However, after treatment with berberrubine, visible morphological changes were observed, including wrinkling of the cell membrane (Fig. 2C), and even rupture and cytoplasmic leakage (Fig. 2D). The severity of cellular damage increased with higher concentrations of berberrubine. These results supported the hypothesis that berberrubine induced oxidative stress and caused membrane damage, thereby exerting its antibacterial effects.

**Fig. 2:**
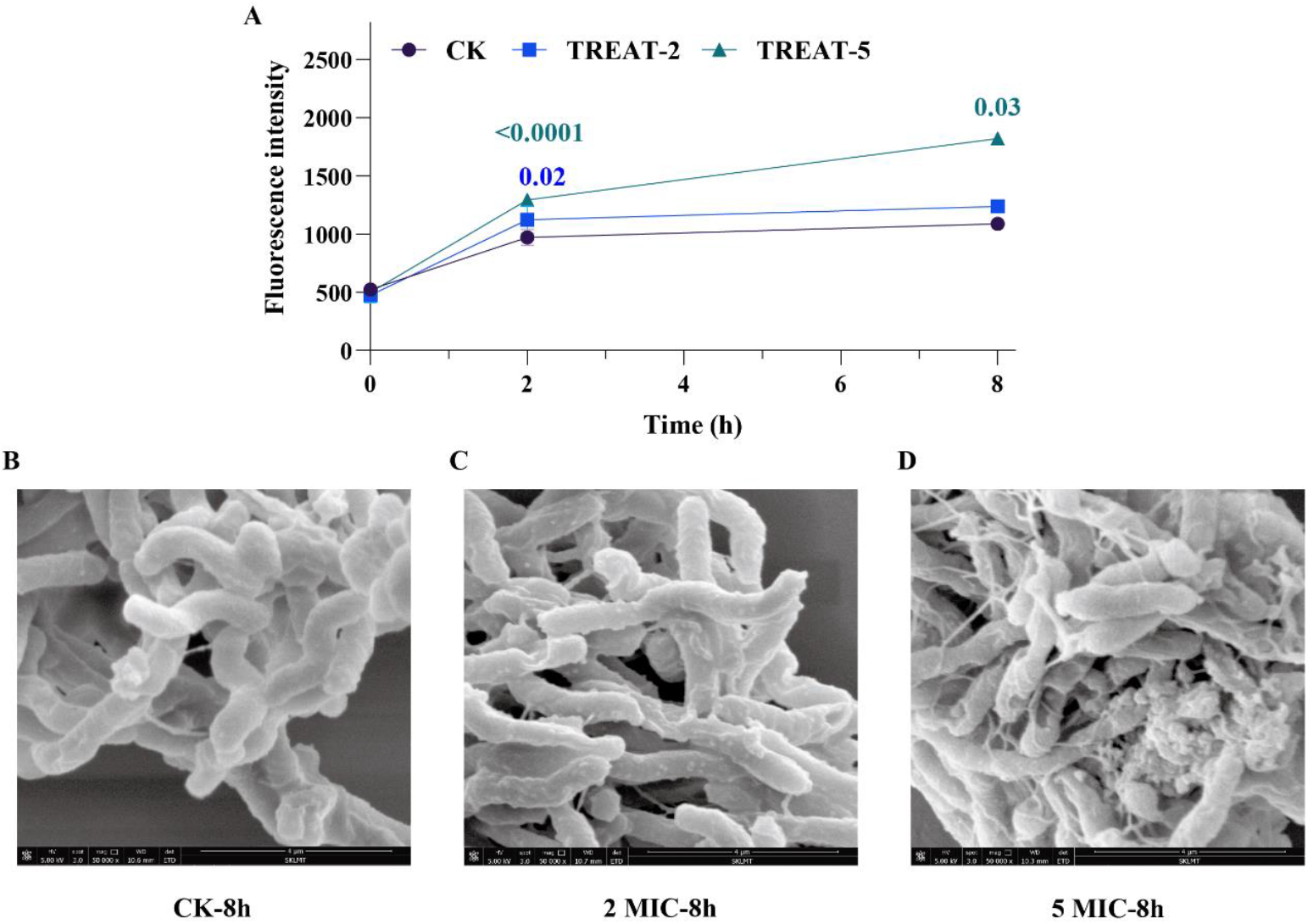
ROS detection and SEM morphology after 2× and 5× MIC treatment of *H. pylori*. (A) Intracellular ROS content detection of *H. pylori*. (B) SEM observation of CK group cell morphology after culture 8h. (C) SEM observation of TREAT group cell morphology after 2× MIC treated 8h. (D) SEM observation of TREAT group cell morphology after 5× MIC treated 8h. Data were shown as the mean±S.E.M. The p value was calculated by Student’s t-test.

### Berberrubine exhibited the same antibacterial mechanism against *Escherichia coli*

To validate the universality of berberrubine’s antibacterial effect, *Escherichia coli*, a representative intestinal opportunistic pathogen, was selected to evaluate its sensitivity to berberrubine and the other 45 natural products previously screened for *H. pylori*. As shown in Table 2, 12 compounds from 5 herbs exhibited antibacterial activity. Similar to *H. pylori*, E. coli was sensitive to all 5 purified compounds and aqueous extracts derived from *Coptidis rhizoma*, with berberrubine showing the strongest antibacterial effect. The MIC of berberrubine against *E. coli* was 0.17 mg/mL.

**Table 2.**
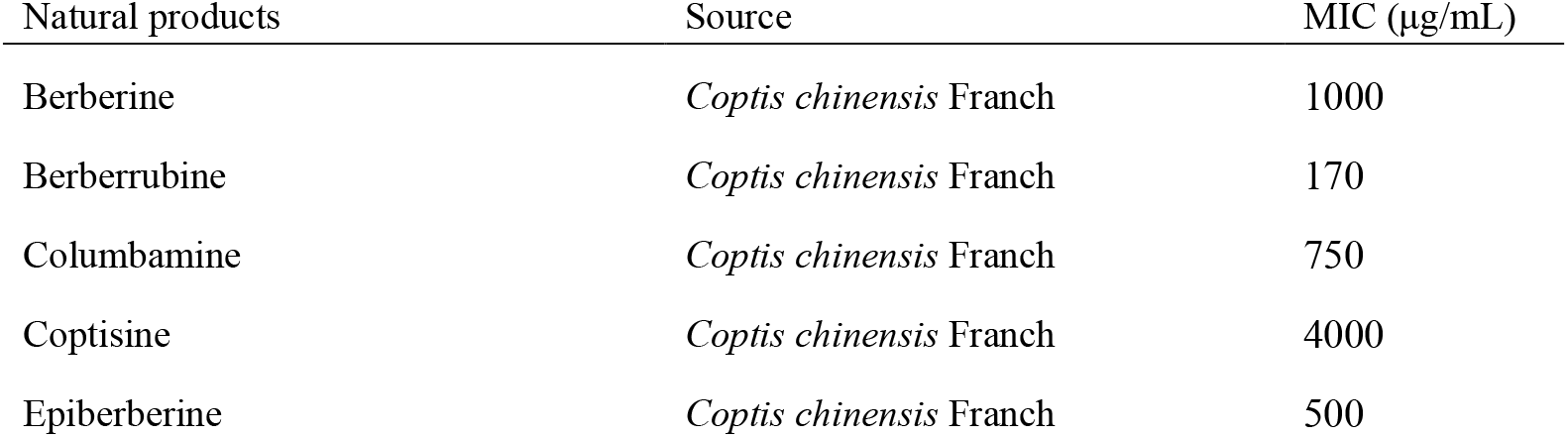
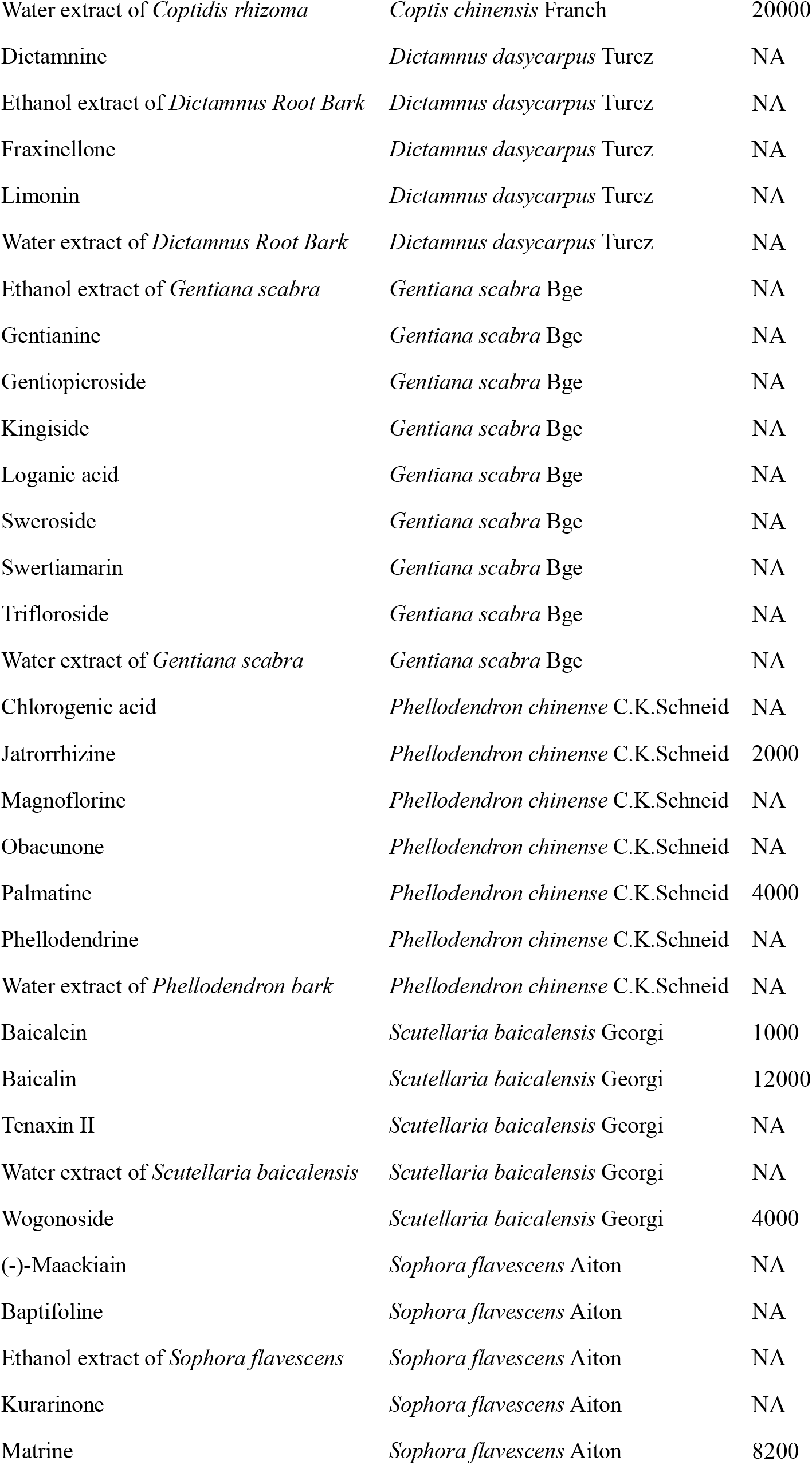
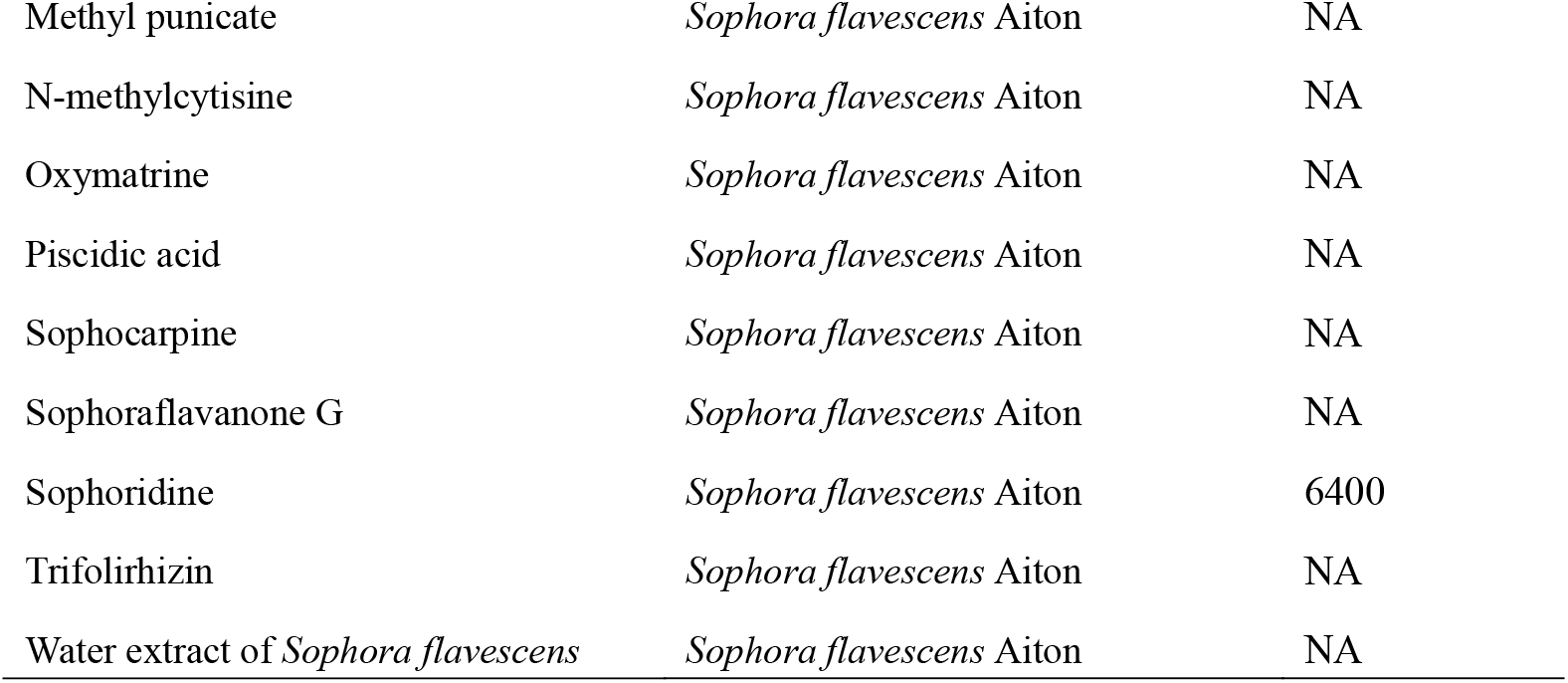
The minimum inhibitory concentration (MIC) of 46 natural extracts against *Escherichia coli*.

To further investigate the similarity of antibacterial mechanism, the genome of *E. coli* was annotated, and the oxidative stress-related genes represented by *sodB* and *fumC*, which were previously identified to increase in *H. pylori*, were also detected in *E. coli*. After treating *E. coli* cells with berberrubine for 2 hours, RT-qPCR analysis revealed significant upregulation of *sodB* and *fumC* (Fig. 3A, B), suggesting that the antibacterial mechanism of berberrubine *in E. coli* might be similar to that in *H. pylori*.

**Fig. 3:**
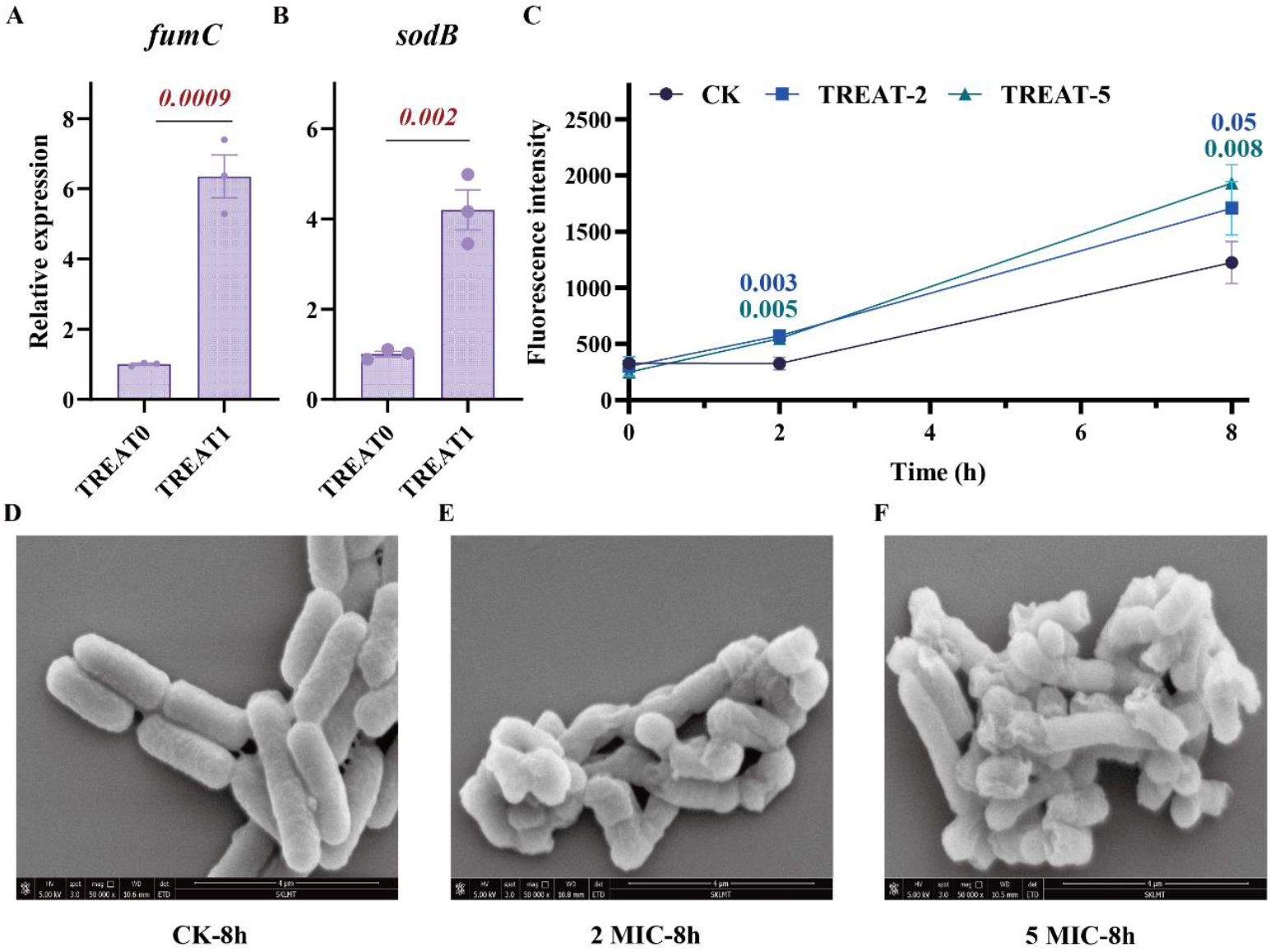
qPCR relative quantification of homologous genes *fumC* and *sodB*, ROS detection and SEM morphology after 2× and 5× MIC treatment of *E. coli*. (A) qPCR relative quantification of *fumC* gene transcription level in *E. coli*. (B) qPCR relative quantification of *sodB* gene transcription level in *E. coli*. Data were shown as the mean±S.E.M. The p value was calculated by Student’s t-test. (C) Intracellular ROS content detection of *E. coli*. (D) SEM observation of CK group cell morphology after culture 8h. (E) SEM observation of TREAT group cell morphology after 2× MIC treated 8h. (F) SEM observation of TREAT group cell morphology after 5× MIC treated 8h.

Furthermore, ROS accumulation was assessed using DCFH-DA after treatment with 2× MIC and 5× MIC concentrations of berberrubine. After 2 hours of treatment, significant increases in intracellular ROS levels were observed in both 2× MIC and 5× MIC treatment groups compared to the control (CK) group (Fig. 3C). Similar to the effect in *H. pylori*, ROS accumulation in *E. coli* was dose-dependent and persisted until 8 hours. Further SEM analysis revealed that, in contrast to the structurally intact cells in the CK group, berberrubine-treated *E. coli* exhibited membrane wrinkling of the cell membrane and cellular damage, with these morphological changes becoming more severe as berberrubine concentrations increased (Fig. 3D-F). These results indicated that, similar to *H. pylori*, berberrubine induced oxidative stress and caused cellular damage in *E. coli*, thereby exerting its antibacterial effect.

## Discussion

*H. pylori* was a major pathogenic bacterium related to chronic gastritis, peptic ulcers, and gastric cancer (9, 10, 35). The standard treatment for *H. pylori* infection relied on antibiotics, with quadruple therapies being widely used (36). However, the increasing resistance to commonly used antibiotics posed significant challenges to successful eradication. Therefore, there was an urgent need to explore alternative treatments, particularly those based on natural products, which have shown promising potential in overcoming antibiotic resistance (12-14). In this study, we screened a range of pure compounds and herbal extracts for their ability to inhibit *H. pylori*.

Previous studies have shown that berberine, from the same family as berberrubine, also has antibacterial effects (28, 29, 37). In the present work, we observed a significant enhancement in the antimicrobial activity of berberine against *H. pylori*, with the MIC decreasing to 25 μg/mL, representing a two-fold improvement over previous reports (29). Additionally, we found that the MIC of berberine against *E. coli* was 1 mg/mL, significantly lower than previously reported values (MIC = 1024-2048 μg/mL) (38). This discrepancy may be attributed to differences in the purity of the compound, which can result from variations in solvent selection, preparation methods, and dissolution conditions. These factors could affect the bioavailability and diffusion of berberine, influencing its antibacterial activity. Alternatively, variations in *H. pylori* and *E. coli* strains and experimental conditions might also contribute to the observed discrepancies in MIC values. Notably, we found that berberrubine exhibited an even stronger antimicrobial effect, with its activity being approximately 2 times higher than that of berberine. Besides, berberrubine also demonstrated a remarkable ability to inhibit the growth of *E. coli*, showing an antimicrobial effect 5 times stronger than that of berberine. Compared to the most commonly used antibiotic therapies for treating *H. pylori*, the MIC of berberrubine (0.011 mg/mL) is even slightly lower than that of clarithromycin (MIC 0.013 mg/mL), levofloxacin (MIC 0.012 mg/mL), and furazolidone (MIC 0.012 mg/mL), indicating that it is a highly promising antibiotic alternative for treating *H. pylori* infections (39). These findings highlight the broader potential of berberrubine as an antimicrobial agent, particularly in combating *H. pylori* and other common pathogens. Based on these promising preliminary results, we plan to further investigate the pharmacokinetics of berberrubine *in vivo*. This will provide critical data for developing it as a novel drug for the treatment of *H. pylori* infections, offering a viable alternative to traditional antibiotics. Given the growing concerns over the misuse of antibiotics and the associated public health risks, our goal is to reduce reliance on antibiotics by providing a safe and effective treatment strategy, thereby mitigating the emergence of antibiotic resistance.

We found that antibacterial mechanism of berberrubine appeared to involve the induction of cellular oxidative stress. We observed significant upregulations of oxidative stress-related genes, represented by *fumC* and *sodB*, a significant increase in cellular ROS level, and a severe cellular damage, after berberrubine treatment. Actually, ROS induced damage to cellular components, including the membrane and DNA, was a well-established mechanism by which antimicrobial agents exert their bactericidal effects (40, 41). Our study also found that berberrubine’s antibacterial mechanism in *E. coli* was similar to that in *H. pylori*, suggesting that the compound might have a broad-spectrum antibacterial effect. These results indicated that berberrubine was a promising candidate for treating a variety of bacterial infections, beyond just *H. pylori*, with potential applications in both gastrointestinal and systemic infections. While our findings are promising, it is important to acknowledge that the clinical application of berberrubine as an antimicrobial agent requires further investigation. The bioavailability, pharmacokinetics, and potential toxicity of berberrubine need to be carefully evaluated in preclinical and clinical studies. Additionally, future research should explore the molecular pathways involved in berberrubine’s antimicrobial action and assess its effectiveness in *in vivo* models.

## Conclusions

In this study, we screened and identified berberrubine as a potent antibacterial drug candidate against both *H. pylori* and *E. coli*, with its mechanism involving the induction of oxidative stress and cellular damage. These findings position berberrubine as a promising candidate for further development as an antimicrobial agent, particularly in the treatment of gastrointestinal and opportunistic infections, in light of the growing challenge of antibiotic resistance.

## Materials and methods

### Natural components preparation

The ethanol and water extracts of several medicinal plants were prepared and analyzed in our laboratory. *Gentiana scabra* Bge. water extract was prepared by soaking the dried roots and rhizomes overnight, followed by reflux extraction three times with 10-fold distilled water (w/v) for 3 h each. The combined extracts were concentrated using rotary evaporation at 60°C and freeze-dried, yielding a crude extract with a 53.5% extraction rate. In addition, the ethanol extract of *Gentiana scabra* Bge. was prepared, and the following compounds were isolated and identified: gentiopicroside, trifloroside, swertiamarin, sweroside, loganic acid, kingiside, and gentianine.

For *Sophora flavescens* Aiton, the dried roots were first soaked for 30 min and subjected to reflux extraction twice with 30-fold distilled water (w/v) for 20 min each time, yielding a water extract with a 25% extraction rate after concentration and freeze-drying. The dried roots were also extracted five times with 3-fold 70% ethanol (w/v) for 1.5 h each time, and the ethanol extract was obtained with a 26% yield after concentration and freeze-drying. The following compounds were isolated and identified: matrine, oxymatrine, sophocarpine, sophoridine, baptifoline, N-methylcytisine, trifolirhizin, kurarinone, (−)-Maackiain, sophoraflavanone G, piscidic acid, and methyl punicate.

For *Dictamnus dasycarpus* Turcz., the dried root barks were pulverized and extracted twice with 10-fold distilled water (w/v) for 2 h each time. The combined extracts were concentrated and freeze-dried, yielding a water extract with a 21.9% extraction rate. The ethanol extract of *Dictamnus dasycarpus* Turcz. was obtained similarly, with a yield of 20.8%. Fraxinellone, limonin, and dictamnine were isolated and identified.

For berberrubine, in a round-bottom flask, 1.01g of berberine hydrochloride and 15ml of N,N-Dimethylformamide (DMF) were added, along with a few boiling stones. The mixture was refluxed under condensation at 190°C for 20 minutes. Then the reaction flask was removed and while hot, 1.5 times the amount of water was added to dilute and to cool the solution. The solution was refrigerated overnight to allow complete crystallization. Finally, perform suction filtration and dry to obtain 0.7747g of red powdery solid - berberrubine, with a yield of 76.7%. Through further purification, berberrubine with a purity of > 95% was obtained.

Lastly, the dried bark of *Phellodendron chinense* C.K.Schneid. was ground into powder, soaked for 30 min in ten volumes of distilled water (w/v), and boiled twice for 2 h per time. The combined filtrates were concentrated and freeze-dried, yielding a water extract with a final extraction rate of 19.2%. Palmatine, jatrorrhizine, obacunone, phellodendrine, magnoflorine, and chlorogenic acid were isolated and identified. *Coptis chinensis* Franch and *Scutellaria baicalensis* Georgi extracts were prepared in our laboratory following previous studies (42, 43). All compounds were identified using HPLC with purities >95%.

### Strains, reagents, and media

The bacterial strains used in this study were *H. pylori* and *E. coli*, which were stored in our laboratory. *H. pylori* was cultured in modified Brain Heart Infusion (BHI) medium, which was supplemented with 3.5 g/L yeast extract, 17.5 g/L peptone, and 10% fetal bovine serum (v/v), while *E. coli* was cultured in MH broth medium. Both BHI and MH broths were purchased from Hopebio, Qingdao, China. 46 natural products were obtained from Tianjin University of Traditional Chinese Medicine, Tianjin, China. The powders were dissolved in 100 μL dimethyl sulfoxide (DMSO) and then diluted with 900 μL of deionized water (ddH_2_O) to a final volume of 1 mL.

### Antimicrobial activity test

The minimum inhibitory concentrations (MICs) of 46 natural products against the tested strains were determined using the Clinical and Laboratory Standards Institute (CLSI) standard broth microdilution method (44). The initial concentrations of the natural products are provided in Table S2, and a total of 11 serial dilutions were performed. The positive control group included a PBS group, with three replicates for each test. The 46 natural products were serially diluted 2-fold in BHI or MH broth in sterile 96-well plates. Each well was then inoculated with 100 μL of bacterial suspension at a concentration of 1 × 10^10^ cells/mL. After co-incubation at 37 °C for 48 hours (for *H. pylori*) or 24 hours (for *E. coli*), the MIC was defined as the lowest concentration at which no noticeable turbidity was observed, by comparing the drug concentration to that in the control wells.

### Transcriptome sample preparation

Based on the previously determined sublethal concentrations of berberrubine against *H. pylori*, 2 sets of samples were selected for transcriptomic analyses to explore the antibacterial mechanisms of berberrubine. The bacteria were inoculated with 1% (v/v), cultured at 37°C until the OD_600_ = 0.6, and then treated as follows: 1) CK: sterile modified BHI without berberrubine; 2) TREAT: sterile modified BHI with 1/2 MIC berberrubine, and then incubated at 37°C for 2 h. After incubation, the cell suspensions were centrifuged at 10,000 rpm for 2 min to remove the culture medium. The cell pellets were immediately frozen with liquid nitrogen and temporarily stored at −80°C before the following RNA extraction.

### Read alignment and normalization of gene expression levels

Raw data (fastq format) were processed using in-house Perl scripts. In this step, clean data (clean reads) were obtained by removing reads containing adapters, poly-N sequences, and low-quality reads. The Q20, Q30, and GC content of the clean data were then calculated. All downstream analyses were based on these high-quality clean data. The reference genome and gene model annotation files were downloaded from the National Center for Biotechnology Information (https://pmc.ncbi.nlm.nih.gov/) (Ref. GCA_007836855.1). The reference genome index was built and paired-end clean reads were aligned to the reference genome using Hisat2 v2.0.5 (45). FeatureCounts (v1.5.0-p3) was used to count the reads mapped to each gene, and FPKM (Fragments Per Kilobase of transcript sequence per Million base pairs sequenced) was calculated based on gene length and read count. Differential expression analysis between conditions was performed using the DESeq2 R package (v1.20.0). Genes with an p-value < 0.05 and the absolute value of log_2_FC > 0 were considered differentially expressed. DEGs functional enrichment was performed using the Gene Ontology (GO) databases. Significantly enriched GO terms were obtained with the criteria of an p-value < 0.05.

### Real-time quantitative reverse transcription-polymerase chain reaction (qRT-PCR)

The harvested cells were collected after being exposed to 1/2 MIC berberrubine. A 5 mL bacterial suspension was centrifuged for RNA extraction. Total RNA was extracted using TRIzol reagent (Vazyme, China). RNA was then reverse transcribed into cDNA using HiScript III All-in-One RT SuperMix Perfect for qPCR (Vazyme, China). The expression of the target genes was measured by real-time qPCR using SupRealQ Purple Universal SYBR qPCR Master Mix (U+) (Vazyme, China) with specific primers (Table S3).

### ROS detection

The cells were centrifuged at 10,000 rpm for 2 min, and the supernatant was discarded. The cell concentration was then adjusted to OD_600_ = 1.0. The cells were incubated with a 10 μM 2′,7′-dichlorofluorescein diacetate (DCFH-DA) solution in the dark for 30 minutes. Unbound DCFH-DA was removed by centrifugation and washed twice with PBS. Fluorescence intensity was measured at excitation and emission wavelengths of 488 nm and 525 nm, respectively, using a multifunctional enzyme reader.

### Scanning electron microscope

2 mL of the culture medium, centrifuged at 8,000 rpm for 5 min, discard the supernatant, and added an equal volume of 2.5% glutaraldehyde solution, incubated at 4°C for 4 h. Washed 3 times with PBS, followed by dehydration through a graded ethanol series: 30%, 50%, 70%, 85%, and 95% ethanol once each, and 100% ethanol twice, with 15-20 min for each step. Replace with tert-butyl alcohol twice, 20 minutes per time.

### Statistical analysis

All data were analyzed using GraphPad Prism 10 software and presented as the mean ± S.E.M. Figures were visualized using Adobe Illustrator. Data comparisons were performed using Student’s t-test.

## Acknowledgments

We thank Sen Wang and Xiangmei Ren from State Key Laboratory of Microbial Technology of Shandong University for the assistance provided in scanning electron microscopy sample preparation and imaging. This research was funded by the National Natural Science Foundation of China (Project No. 82030116).

## Data availability

Data is deposited in National Microbiology Data Center (NMDC) with accession numbers NMDC10019606 (https://nmdc.cn/resource/genomics/project/detail/NMDC10019606).

## Contributions

SJ L, C L and F Q conceived and designed research. MZ J conducted experiments. CY W, Y J and X W provided experiment bacteria. MZ J analyzed data and wrote the manuscript. All authors read and approved the manuscript.

## Corresponding author

Correspondence to Feng Qiu, Chang Liu and Shuang-Jiang Liu.

## Ethics declarations

## Ethical approval

No ethical approval.

